# Position-specific evolution in transcription factor binding sites, and a fast likelihood calculation for the F81 model

**DOI:** 10.1101/2023.07.23.550199

**Authors:** Pavitra Selvakumar, Rahul Siddharthan

**Affiliations:** The Institute of Mathematical Sciences, Chennai, India; Homi Bhabha National Institute, Mumbai, India

## Abstract

Transcription factor binding sites (TFBS), like other DNA sequence, evolve via mutation and selection relating to their function. Models of nucleotide evolution describe DNA evolution via single-nucleotide mutation. A stationary vector of such a model is the long-term distribution of nucleotides, unchanging under the model. Neutrally evolving sites may have uniform stationary vectors, but one expects that sites within a TFBS instead have stationary vectors reflective of the fitness of various nucleotides at those positions. We introduce “position-specific stationary vectors” (PSSVs), the collection of stationary vectors at each site in a TFBS locus, analogous to the position weight matrix (PWM) commonly used to describe TFBS. We infer PSSVs for human TFs using two evolutionary models (Felsenstein 1981 and Hasegawa-Kishino-Yano 1985). We find that PSSVs reflect the nucleotide distribution from PWMs, but with reduced specificity. We infer ancestral nucleotide distributions at individual positions and calculate “conditional PSSVs” conditioned on specific choices of majority ancestral nucleotide. We find that certain ancestral nucleotides exert a strong evolutionary pressure on neighbouring sequence while others have a negligible effect. Finally, we present a fast likelihood calculation for the F81 model on moderate-sized trees that makes this approach feasible for large-scale studies along these lines.

## Introduction

Transcription factor binding sites (TFBS) are short regions in DNA where regulatory proteins bind, typically by recognizing a short, conserved “motif”. These motifs are not exact strings and are commonly represented by “position weight matrices” (PWMs) [1], whose columns give position-specific nucleotide distributions: for a motif of length *L*, a PWM *W*_*nα*_ is a *L ×* 4 matrix giving the probability, if a sequence of length *L* is a binding sequence, that the nucleotide at position *n* is *α*.

The PWM representation assumes that different positions within a motif (different columns of the matrix) have independent distributions of nucleotides. However, positional dependencies are known to occur in binding sites of many TFs. Extensions to the basic PWM format have been proposed to account for positional dependencies [2, 3, 4, 5]. Nevertheless, PWMs persist as the dominant motif representation format, because of ease of calculation and because they are easily visualized via sequence logos [6].

In this work we ask: in the course of evolution, what are the selective pressures on TFBS? Are these pressures position-independent, or do nucleotide preferences at one position affect evolution at other positions?

Several models of DNA evolution have been developed in the past decades, briefly reviewed in Methods. Neutral models of DNA evolution assume that individual nucleotides mutate at a rate given by a rate matrix, from which can be calculated transition probabilities at finite time, and long-term distributions at individual loci. Symmetric rate matrices, such as in the Jukes-Cantor [7] and Kimura [8] models, yield a uniform stationary distribution of nucleotides under neutral evolution. Commonly, selection is treated as an additional process affecting nucleotide evolution and distribution. In the context of site-specific evolution, the Halpern-Bruno model, developed to describe protein codon evolution [9], was modified for TFBS evolution by Moses *et al*. [10]. The assumptions are that TFBS are under purifying selection (where deleterious mutations are removed from the population), and that time to fixation is small compared to time between fixations (“weak mutation model”). This is further explored in Discussion.

Here we take an alternative approach: we consider evolution via a rate matrix that yields a non-uniform stationary distribution, and seek to infer that stationary distribution (which is indicative of fitness under purifying selection) via an evolutionary model. The earliest such model, and the simplest, was Felsenstein’s 1981 model (F81) [11]. (In the same paper, Felsenstein supplied his now-standard recursive “pruning” algorithm for likelihood calculation). We also use the Hasegawa-Kishino-Yano 1985 (HKY85) [12] model, which yields similar results on this problem.

Our approach is as follows: we use a standard tree topology from the literature (for primates, and for mammals); we obtain instances of motifs (with 20bp flanking sequence) in human, and orthologous motifs (without indels) in other species, for various TFs; we estimate the tree’s branch lengths using the least informative positions in these sequences, which we assume to be neutrally evolving (stationary vector equal to background probability distribution of nucleotides in human); and then, given this complete tree with topology and branch lengths, we estimate “position-specific stationary vectors” (PSSVs) at all positions within the motif.

These PSSVs are the long-term stationary distribution of sites evolving under our evolutionary model and are obtained via maximum likelihood, as described in Methods. They can be seen as combining mutation and selection information. Our motivation is to study how they compare to a PWM, how they may vary across different factors, and how they may vary if we change the set of species studied (eg, from primates to a larger set of mammals).

Our methodology enables inference of ancestral nucleotide probabilities at each locus. This allows us to study compensatory mutations via “conditional PSSVs”, by identifying a subset of sites where the ancestral nucleotide at a given position is, say, G with probability *>* 0.5, and then studying the effect of the PSSV at other positions within this subset of sites.

En route, we present an algorithm for calculating likelihoods in the F81 model, which, for moderate-sized trees, is much faster than Felsenstein’s pruning algorithm. As described in Methods, the idea is to reduce an arbitrary tree into a sum of products of “star trees”, where we define a star tree as a tree all of whose leaves are directly connected to one ancestral node. The speedup comes partly from reducing the complexity in alphabet size Σ from *O*(Σ^2^) (pruning algorithm) to *O*(Σ), but mainly from removing the recursive traversal of the tree which otherwise needs to be done at every locus.

## Materials and Methods

### General approach

Given a collection of orthologous sequences from a set of species, and given a phylogenetic tree linking those species, we can calculate, via standard methods described below, the likelihood of those sequences arising via an evolutionary model from a common ancestor. We take the topology of the tree from the literature; these are shown in Figure 1 (A) and (B). We focus on TFBS in human (including flanking sequence), with gapless orthologous sequence in other species. We assume that uninformative positions have a uniform stationary vector (equal distribution of probabilities under evolution) and estimate the branch lengths (the evolutionary time *t* between neighbouring nodes of the tree) from these positions, via maximum likelihood. We use two evolutionary models, the Felsenstein 1981 (F81) below, and the Hasegawa-Kishino-Yano 1985 (HKY85) model, described in the appendix, that are capable of reproducing arbitrary stationary distributions; having learned the tree branch lengths from uninformative positions, we then learn the site-specific stationary vectors that would be best explain the data at other positions, again via maximum likelihood.

**Figure 1:**
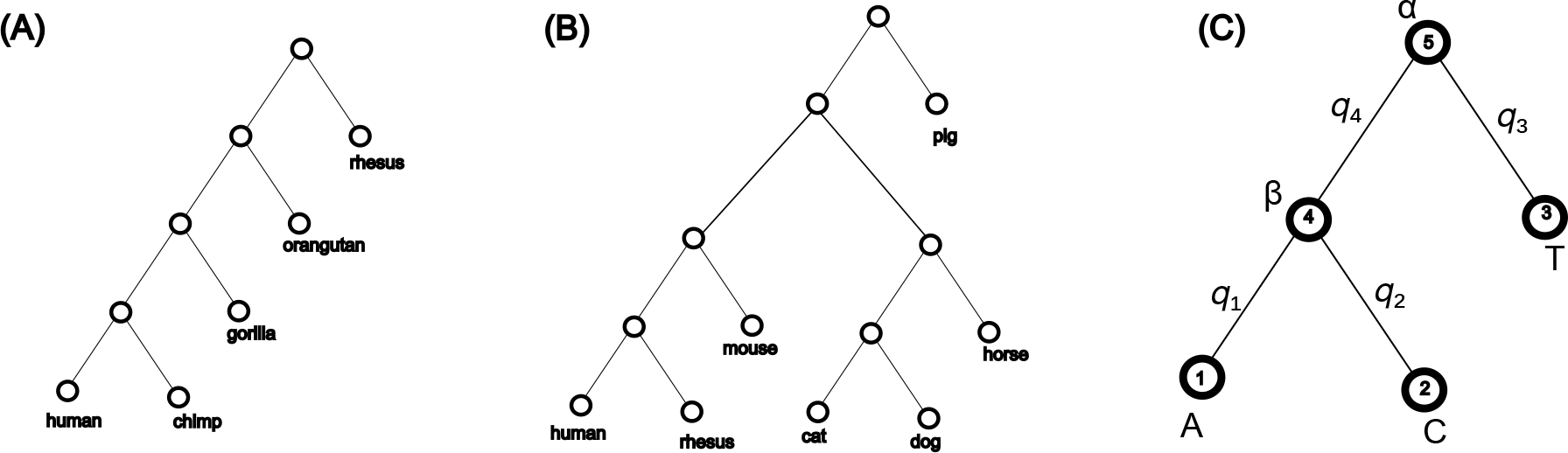
(a) Primate tree topology used in this work (b) Mammal tree topology used in this work (branch lengths are calculated and not shown in this figure) (c) Sample tree with proximity labels on branches

Necessary background and notation is established in the appendix. Let *T*_*αβ*_(*t, μ*) be the probability, given ancestor *β*, a mutation rate *μ* and an evolutionary time *t*, that one observes *α* at the same locus. We choose units of time such that *μ* = 1, and define the “proximity” *q* ≡ *e*^−*t*^ [13]. In terms of this, the F81 model is:

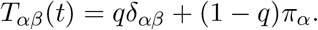

Further discussion is in the appendix.

### Fast likelihood calculation for F81 model

The F81 transition matrix has a peculiar interpretation that allows us to greatly speed up likelihood calculations for moderate-sized trees, compared to the Felsenstein pruning algorithm (described in the appendix). First, we generalize trees beyond binary trees, so that nodes can have more than two children. We define a “star topology” as a tree topology where all nodes, except the root, are leaves (i.e. every leaf is directly attached to the root). It turns out that, under the F81 model, *every tree is equivalent to a sum of products of star-topology trees*.

We illustrate with fig. 1 (c), where we had (in equation 4)

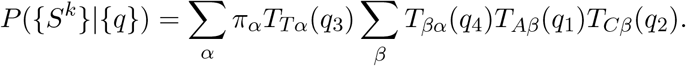

Consider the sum over *β*. The node corresponding to that sum is what we call a “star node”, i.e. a node all of whose children are leaves. We plug in the explicit expression *T*_*βα*_ = *q*_4_*δ*_*αβ*_ + (1 − *q*_4_)*π*_*β*_, to obtain

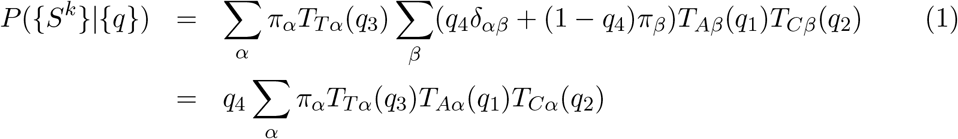

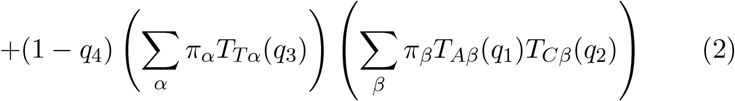

Basically, with probability *q*_4_ the node *β* is unmutated from its parent, so its leaves can be directly joined to its parent (the first term in eq. 2). And with probability 1 − *q*_4_ it is mutated but is now completely independent of its parent *α* and its likelihood now merely multiplies the rest of the tree.

For this simple tree, this is all, but we note that

- Every tree that is not a star tree has at least one star node
- A procedure equivalent to that in equation 2 can be applied to that node, by summing over its unknown nucleotide, and it has the same effect.

Suppose the original tree is *T*, and the subtree starting at the star node is *T*_*s*_. Let the star node be *i* and its parent be *j*. The full likelihood for the tree will contain the following for this node:

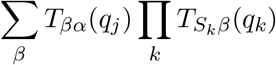

where *k* are the children of *j*, which are all leaves, *α* is the nucleotide at *j*’s parent, and as always, *q*_*i*_ is the proximity of node *i* to its parent. Using

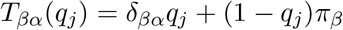

the original likelihood expression can be split into two terms, in one of which (with weight *q*_*j*_) *T*_*s*_ is merged with its parent to make a merged tree *T*_*m*_, and in the other (with weight 1 −*q*_*j*_) *T*_*s*_ is removed from its parent and separated out, yielding *T* = *q*_*j*_*T*_*m*_ +(1 −*q*_*j*_)*T*_*p*_*T*_*s*_.

This can be done iteratively until the tree is entirely a sum of product of star trees:

1. Initialize a list of lists of trees, [*L*_1_, *L*_2_, *L*_3_ …], initially with a single tree list [L] with a single element *L* = [*T*] (the original tree)
2. For each list of trees *L* = [*T, T* ^*′*^, *T* ^*′′*^ …]
  a. The first element *T* is the “main” tree, the rest are multipliers
  b. While *T* has non-star nodes
    i. identify a star node in *T*, whose proximity to its parent we call *q*. This can be done in worst-case linear time in number of leaves, by depth-first search.
    ii. create three new trees from *T* : *T*_*m*_ (where star node’s leaves are merged into parent), *T*_*p*_ (where star node and its leaves are removed from parent), *T*_*s*_ (the star node that has been removed).
    iii. Replace original *T* with *T*_*m*_ with a weight *q*; and append a new element to the list consisting of [*T*_*p*_, *T*_*s*_, …] with weight 1 − *q*, where … indicates that all previous multipliers are retained
    iv. All multipliers are star nodes, so the only possible non-star trees are the first elements in the tree lists
  c. Repeat until no non-star trees remain in this list

Under this process, the list of lists of trees may, for example, evolve as follows:

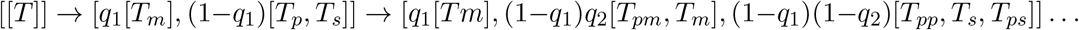

where *T*_*pm*_ indicates a previously pruned tree that has been merged, and so on. This need be done only once for each tree, and then the “sum-of-stars” tree list can be used for fast likelihood calculations on that tree.

For every star node, two new trees are created. In a typical tree, the number of star nodes is proportional to the number of leaves (in a perfectly balanced tree it is half the number of leaves). Therefore, the number of terms in the “sum of trees” grows rapidly with the number of leaves. If the number of leaves is much greater than about 12, this method is not usable (see Results).

We tested our implementation on synthetic and real data and the likelihood calculated is in agreement with the full Felsenstein method,to machine precision. The code is available on the notebooks in the github link listed in “Data availability”.

### Position-specific stationary vectors for TFBS

Our approach is to calculate the *q*’s first (and, for HKY85, *κ*), which we assume are not site-specific, using unconserved/uninformative sites (which we assume are evolving more-or-less neutrally) and assuming a stationary vector *π*_*A*_ = *π*_*T*_ = 0.3, *π*_*C*_ = *π*_*G*_ = 0.2 (the approximate nucleotide distribution in mammals), by maximising the likelihood of these neutral positions. Having fixed the *q*’s and *κ*, we calculate the site-specific 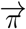’s within TFBS by, again, numerically maximising the likelihood of the observed nucleotides at those positions.

The calculation of *q*’s and *κ* in this work was done, using a specific uninformative position in the neighbourhood of one motif (combining all sites for that motif); the same *q*’s were used for all further calculations in that species. In principle, the *q*’s can also be calculated from other neutral loci, such as synonymous sites in coding sequence (correcting for codon bias).

### Ancestral weight vectors and conditional PSSVs

It is of interest to infer the likely ancestral nucleotide at every locus, and to examine if the ancestral nucleotide has an effect on nucleotides at other positions. We define a “conditional PSSV”, conditioned on a locus *i* and an ancestral nucleotide *α*, as a PSSV trained on a subset of sequences where the ancestral nucleotide at *i* has a probability *>* 0.5 of being *α*.

At every locus, we have a collection of leaves *L*. Let the ancestral node be denoted by *X*. At a locus *i*, the probability of the ancestor *X* having nucleotide *α* can be written as

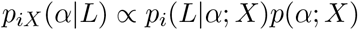

We take *p*(*α*; *X*) (the prior for *α* at *X*) to be *b*_*α*_ (the background probability for *α*, =0.3 for A, T, 0.2 for G, C) and *p*_*i*_(*L* | *α*; *X*) is the likelihood of the leaves at that locus given *α* at the root. This likelihood can be calculated by the fast “sum of stars” method for F81, or by the standard Felsenstein algorithm for other models.

### Data used

ChIP-seq data was obtained from Gene Transcription Regulation Database [14, 15] (GTRD). For each TF, TFBS within chip-seq peaks were predicted using FIMO [16], with the default threshold *p* = 0.0001, with motif models (PWMs) from HOCOMOCO [17]. 20bp flanking sequence was added to each motif, in order to examine possible selection effects on flanking sequence. UCSC LiftOver (command line tool from https://genome.ucsc.edu)was used to find orthologous regions in other organisms. Only gapless liftOver matches were considered. The organisms and assemblies used were:

#### Primates

human (hg38), chimpanzee (PanTro6), gorilla (GorGor6), orangutan (PonAbe3), rhesus monkey (RheMac10)

#### Mammals

human (hg38), rhesus monkey (RheMac10), mouse (Mm39), cat (FelCat9), dog (CanFam6), pig (SusScr11), horse (EquCab3)

The human and orthologous TFBS sequence were stored in a custom file format based on the FASTA format. Processed data for the factors presented here, and other factors, are available in the github link cited in “Data availability”.

### Scrambled PWMs and random genomic sequences

Three scrambled CTCF motifs were considered, with PWMs consisting of the same columns as in the HOCOMOCO motif in a different order. The motif orders were (8, 12, 18, 2, 15, 17, 6, 4, 11, 5, 16, 19, 14, 13, 9, 1, 3, 10, 7), (11, 5, 4, 9, 2, 16, 10, 12, 14, 7, 6, 8, 18, 19, 3, 13, 15, 17, 1), and and (13, 16, 9, 5, 7, 6, 4, 11, 1, 2, 12, 17, 18, 8, 10, 15, 3, 14, 19) where numbers indicate the position of that column in the original motif.

In addition to finding motif instances of these in ChIP-seq peaks, we used FIMO to detect instances of these on 50,000,000 randomly-selected 150bp regions from the complete human genome (excluding the Y and M chromosomes). The regions were uniformly selected (instances in each chromosome proportional to its length) without regard to genome annotation. For this, FIMO was run with the parameter `--thresh 0.000005` (*p* = 5 *×* 10^−6^) rather than the default of *p* = 10^−4^, to obtain more informative PWMs from these motif instances, comparable to the CTCF.

For comparing the “loss of information” of PSSVs compared to PWMs, we calculated, in each case, the information score of the PWM and the PSSV as

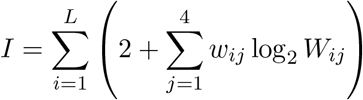

(this is the height of each column in the logo representation, measured in bits), where *i* runs over motif positions, *j* over nucleotides, and *W* is either the PWM or the PSSV matrix. The loss of information for each motif in each dataset is (*I*_PWM_ − *I*_PSSV_)*/I*_PWM_ where *I*_PWM_ and *I*_PSSV_ are respectively the information content of the PWM and PSSV associated with that motif in that dataset.

## Results

### The sum-of-stars F81 likelihood algorithm shows significant speedup for moderate-sized trees

Figure 2 shows the speed of our implementation of the Felsenstein pruning algorithm, compared with our “fast-star” algorithm, for the F81 model on trees of varying sizes and with sequences of varying length. For trees with ≲ 10 leaves we obtain a significant speedup.

**Figure 2:**
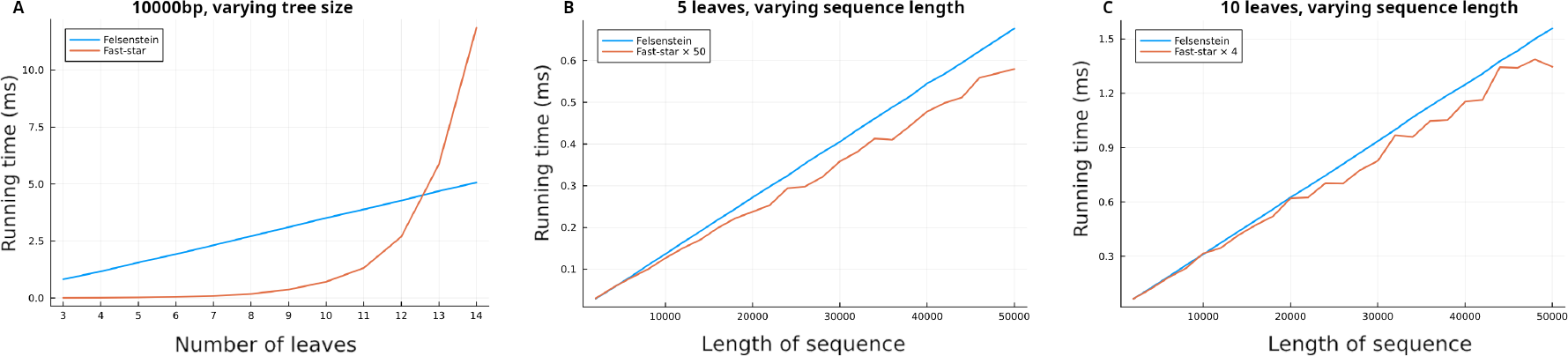
The Felsenstein pruning algorithm’s running time, compared to our “faststar” algorithm. As a function of tree size, we achieve a speedup of *>* 50*×* over the pruning algorithm for trees with 5 leaves, and *>* 4*×* for trees with 10 leaves, over a range of sequence lengths. For larger trees our performance deteriorates.

Further speedup will be possible: our current implementation of fast-star still uses a tree data structure, but star trees can be represented using matrices, which will greatly improve cache performance.

### PSSVs resemble PWMs, but more weakly

Figure 3 shows PWMs, PSSVs and conditional PSSVs (described below) for five representative TFs: CTCF, GABPA, REST, TCF12, USF2. More examples are available via notebooks on github (see Data availability). The PWMs are nucleotide counts within one species (human). PSSVs, position-specific stationary vectors on individual positions derived from an evolutionary model, closely reflect the pattern of the PWM, but are weaker (only slightly weaker for GABPA, but significantly weaker for TCF12, perhaps indicating relative turnover rates of these binding sites).

**Figure 3:**
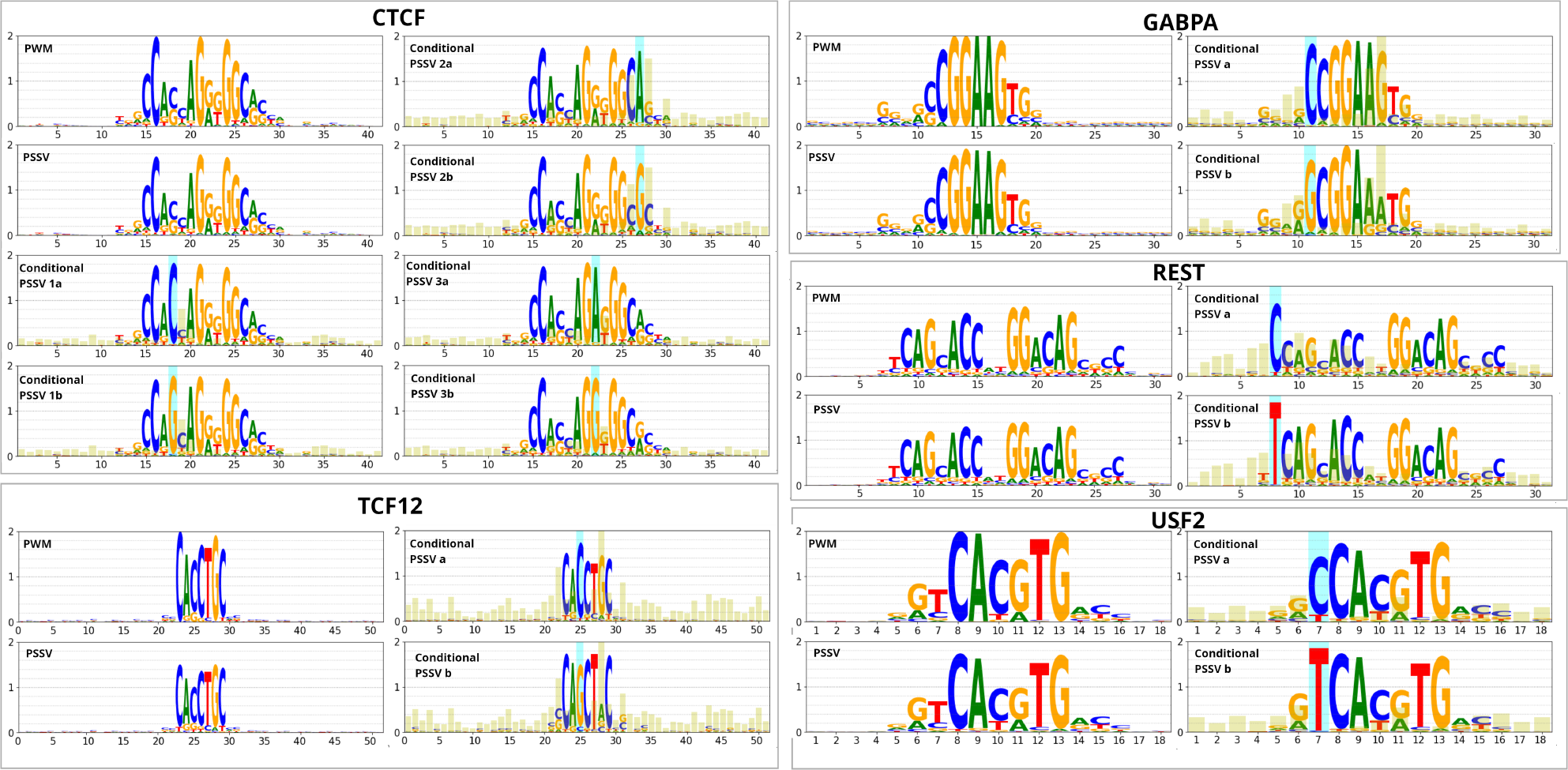
PWMs, PSSVs and conditional PSSVs for five transcription factors, calculated from primate data. The *y* axis in all cases is the number of bits in the information content of the log, ranging from 0 to 2. In conditional PSSVs, the blue highlight is the ancestral nucleotide upon which the PSSV is conditioned, and the yellow highlights show the Jensen-Shannon divergence between the two different conditional PSSVs (JSD*×*5 is plotted for clarity).

One can ask whether this is simply an effect of turnover of sites. Figure 4 compares PWMs constructed from human alone, as in figure 3; PWMs constructed from five primates (human sites plus orthologous sites from four other primates); and PSSVs. The five-primate PWMs are all a little weaker than the human-only PWM, suggesting some level of site turnover, but the PSSV is weaker still, particularly in the case of TCF12.

**Figure 4:**
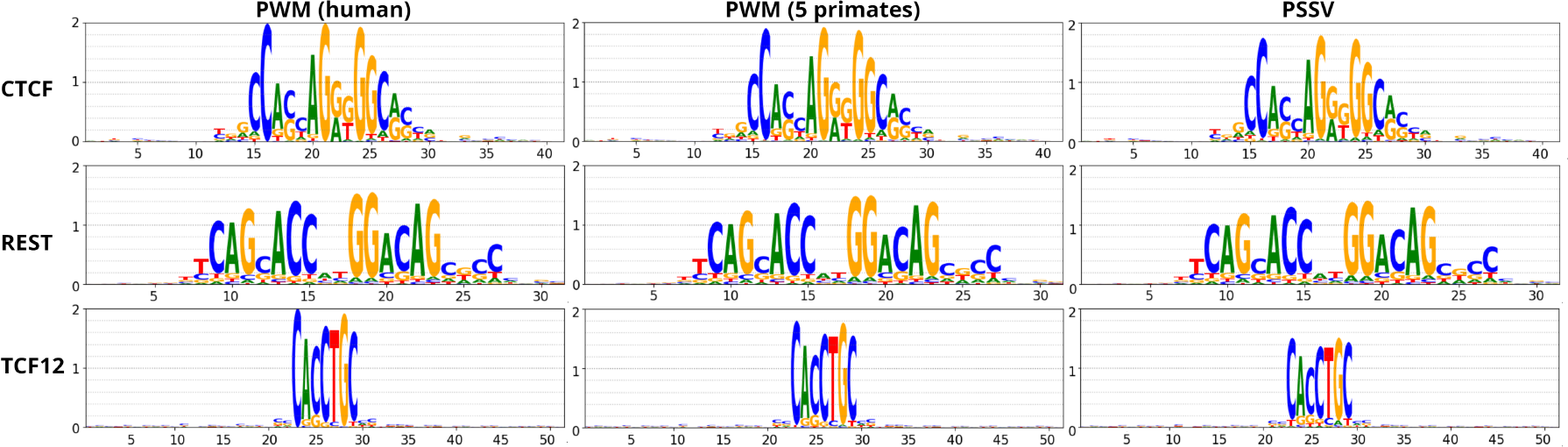
A PWM constructed from human + orthologous sequence from 4 other primates is weaker than a human-only PWM, suggesting some turnover of TFBS; the PSSV is weaker than the 5-primate PWM.

Figure 8 shows further weakening for PSSVs when more distant mammals are included.

### Scrambled PWMs, and random sites from full genome, have less-informative PSSVs

As a control we predicted sites for three examples of scrambled CTCF motif instances (the same core motif columns in a different order), that is, motif instances predicted in the genuine CTCF peaks for these scrambled motifs.

As a further control we predicted sites, for both the genuine CTCF motif and the three scrambled motifs, in random 150bp sequence selected from throughout the human genome. Sites from each chromosome were selected in proportion to its length; the chromatin annotation was not considered. The resulting PWMs and PSSVs, for all these cases, are in figure 5. PSSVs for scrambled motifs, and in the random genomic sites, are visibly reduced in height compared to the CTCF PSSV from ChIP-seq peaks.

We quantify this using loss of information content (see Methods) in fig 6. While the CTCF PSSV from the ChIP-seq peaks has *<* 10% information loss, the PSSV for scrambled PWM motif instances in the ChIP-seq peaks show ≈ 15%–18% information loss, as does the PSSV for CTCF motif instances in random genomic windows. Scrambled PWM motif instances in random genomic sites show ≈ 23%–25% information loss.

**Figure 5:**
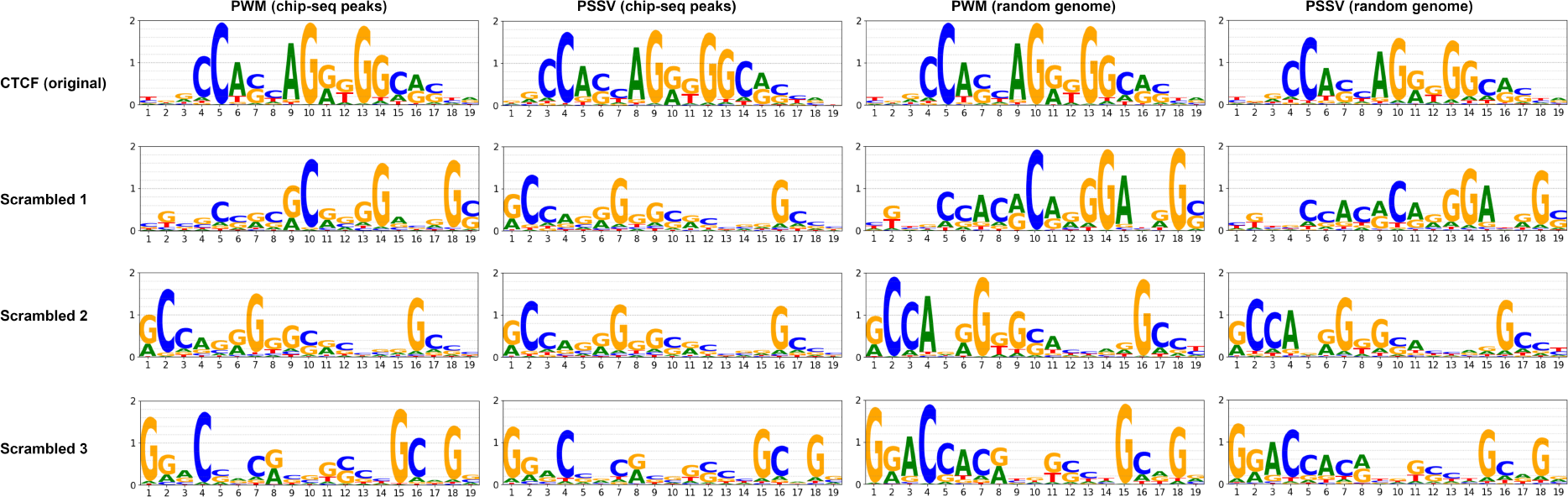
PWMs and PSSVs for the genuine CTCF motif and three scrambled versions, from motif instances found in ChIP-seq peaks for CTCF, and from motif instance found randomly in the genome.

**Figure 6:**
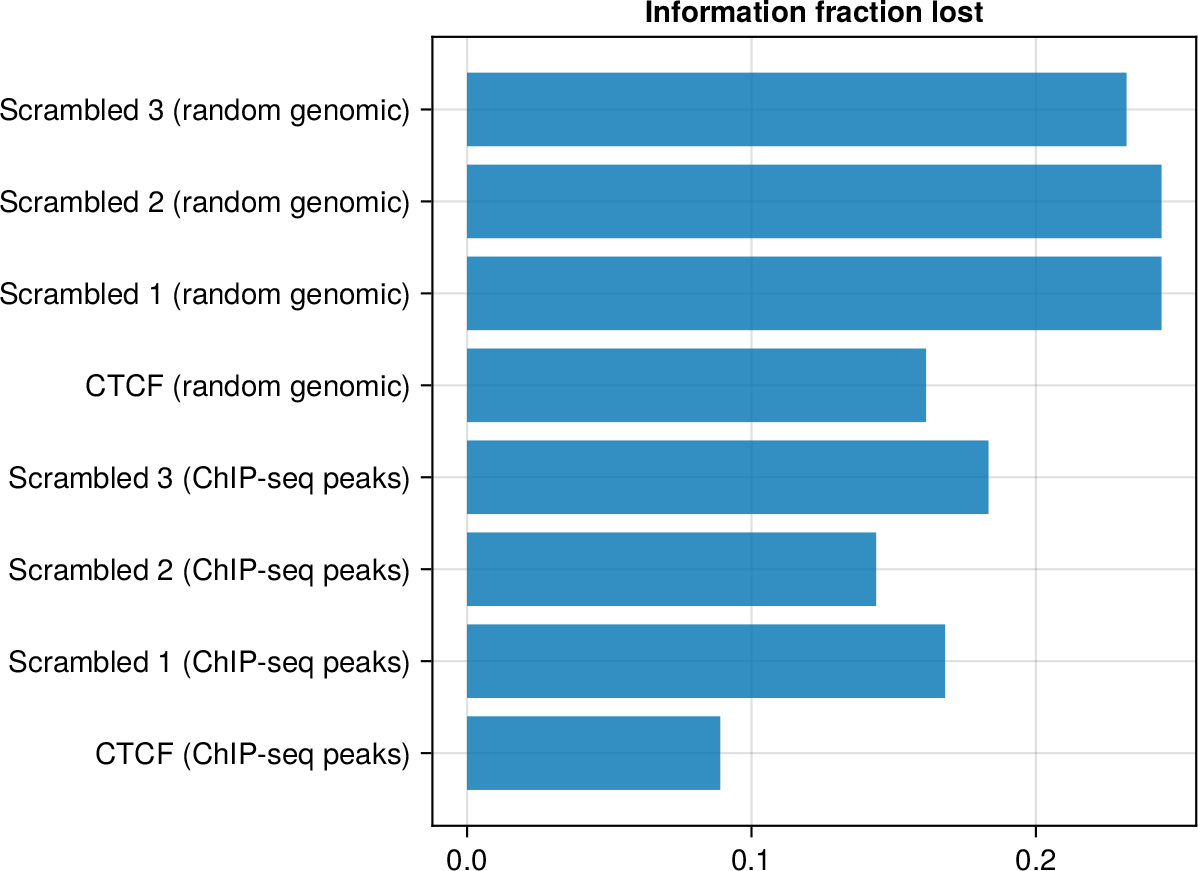
Loss in information for PSSVs compared to PWMs, for CTCF and three scrambled version, in ChIP-seq peaks and in random genomic sequence.

This is consistent with the idea that ChIP-seq regions in general are under selection and, within them, genuine motif instances are under stronger selection, while there is weaker selection in the genome as a whole. We further explore this in Discussion.

### Multiple TFBS exhibit positional correlations based on ancestral nucleotide selections

Figure 3 shows the effect of conditioning certain nucleotides ancestrally on the resulting PSSV, that is, it shows PSSVs for subsets of sites where the inferred ancestral nucleotide is a particular nucleotide *α* with a probability ≥ 0.5. Blue highlights indicate the position of ancestral conditioning, and the yellow highlights are proportional to the Jensen-Shannon divergence between the stationary vectors in the two conditioned sets. CTCF is a widely-studied DNA-binding protein that plays roles in transcriptional regulation, insulator function, and chromatin remodelling [18]. Its binding sequence has long been recognized to have significant positional correlations [19]. We observe here that an ancestral preference for A at position 27 causes a preference of C at position 26 and G at position 28, while an ancestral preference of G at 27 causes a weaker preference of C at 26 but a stronger preference of C at 28. (Eggeling *et al*.[19] had noted a high mutual information between our position 27 and neighbouring positions.) In contrast, an ancestral preference of C or G at position 18 appears to show no obvious change in the PSSV at other positions. (Eggeling *et al*.[19] observe a low mutual information for that position.) An ancestral preference of A or G at position 22 shows weak effects at positions 17 and 26. But also, positions 3 and 5 suggest signs of a motif, previously called the M2 motif [20], that has previously been noted to occur in some CTCF instances; it appears an A at 22 is associated with a slightly stronger M2 motif. We will investigate CTCF further in future work.

Other factors shown in figure 3 show similar positional correlations. Of interest: REST has a dimeric motif, but a choice of C in position 8 seems to suppress the first half of the dimer. And TCF12’s binding sites have CG-rich sequence signatures extending well beyond the core motif, which also reflect in the PSSV, indicating selective pressure in an extended region around the core site. An ancestral choice of G at position 25 appears to strengthen the PSSV overall, including these peripheral CG signatures, but greatly weaken position 28.

Figure 7 shows a similar analysis for the three scrambled CTCF motifs we studied. There is less conditional dependency evident here, but some sign of a dependency in CG dinucleotides, which may be a result of methylation and deamination. Specifically, while the highest JS divergence in the genuine CTCF motif is 0.30 (for position 28 in conditional PSSV 2a/2b, figure 3), the highest JS divergence observed in the three scrambled motifs is 0.11, 0.13 and 0.16 at positions 9, 5, 13 respectively.

**Figure 7:**
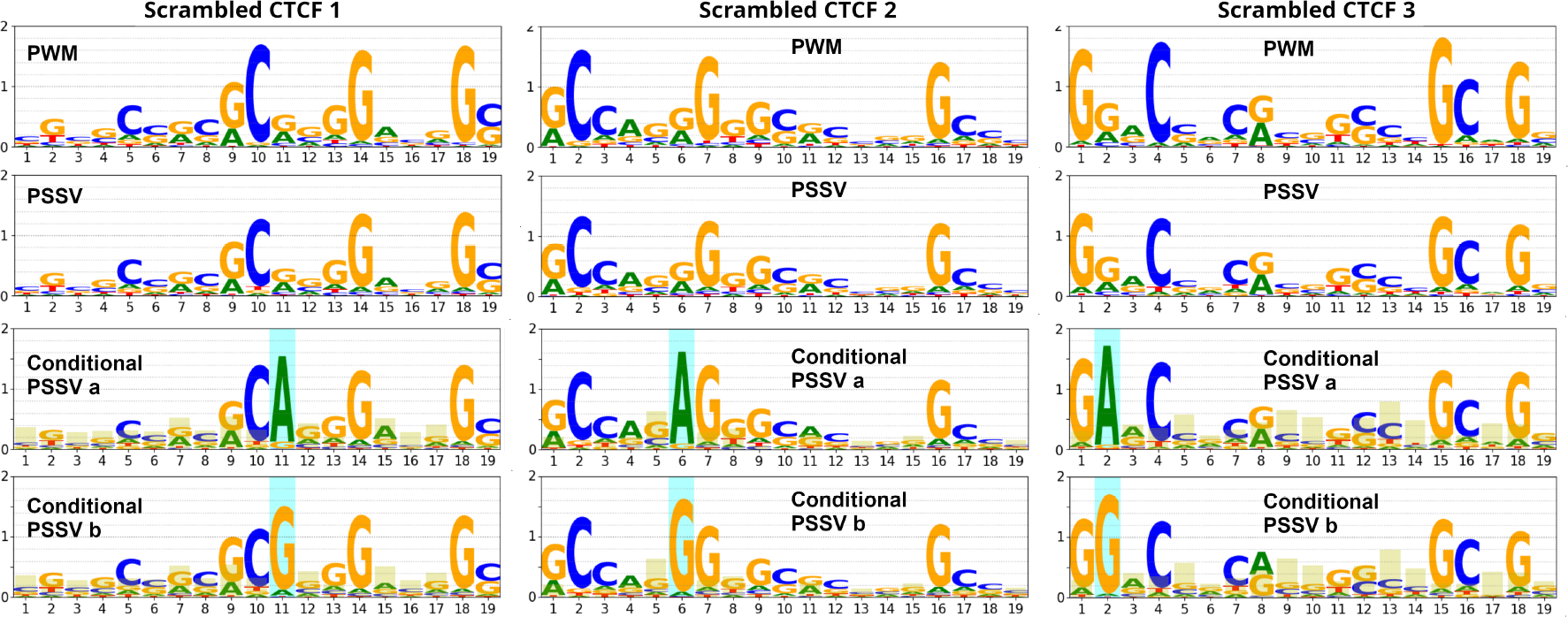
PWMs, PSSVs and conditional PSSVs are shown for site matches for three scrambled versions of the CTCF motif. All are conditioned on the position corresponding to the one highlighted in figure 3, conditional PSSVs 1a and 1b.

We picked positions to condition on, that, in their PWMs, had two dominant nucleotides. An exhaustive conditioning study for some factors (including CTCF) will be the subject of future work.

### The F81 and HKY85 models give similar PSSVs

Figure 8 compares PSSVs obtained from the HKY85 model on five primates, the F81 model on the same 5 primates, and the F81 model on 7 mammals. All are similar, though the F81 mammal PSSVs are weaker.

**Figure 8:**
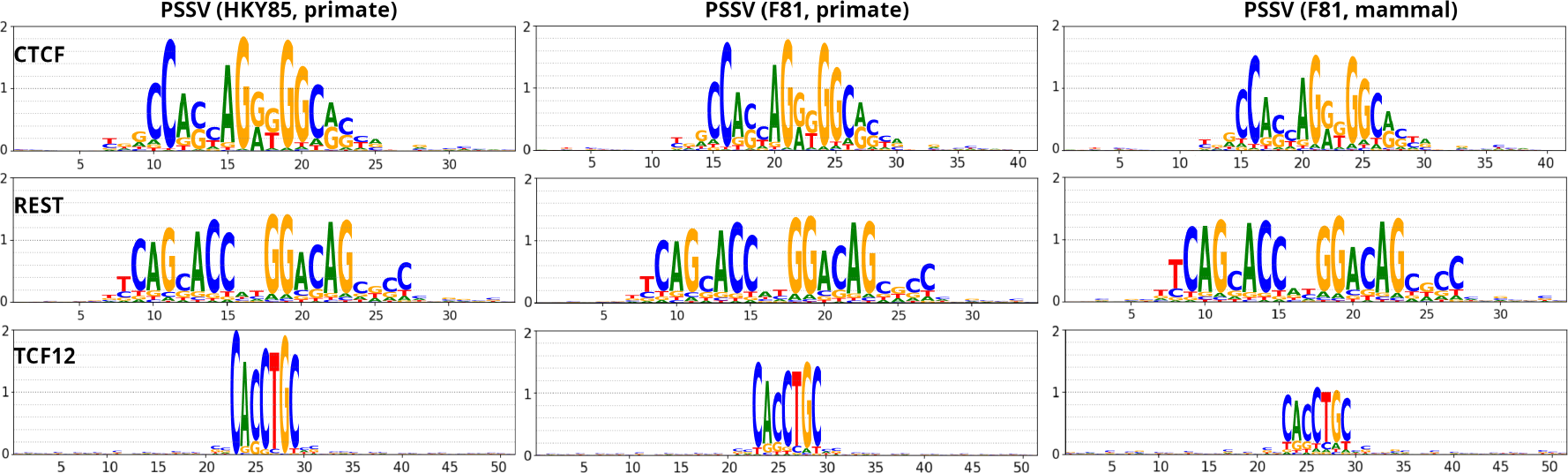
Comparison of PSSVs obtained from the HKY85 model on five primates, the F81 model on primates, and the F81 model on seven mammal species.

## Discussion

TFBS evolution has previously been studied from various points of view. Gerland and Hwa [21] presented a model of molecular evolution that includes genotype, phenotype and fitness, focussing on the fitness advantag to maintain a binding sequence and the evolution of multiple sites in a regulatory region. Similarly, Sengupta *et al*. [22] considered, theoretically, the tradeoff between TFBS specificity and robustness to mutation. Bais *et al*. [23] considered TFBS evolution in the context of annotated genome alignments; they consider both the F81 model and the Halpern-Bruno model. Mustonen *et al*. [24] considered a biophysical model for yeast TFBS evolution, where positional correlations emerge as a consequence of maintaining energy-dependent fitness (via compensatory mutations).

Most similarly to our work, Moses *et al*. [10] discuss position-specific evolution in TFBS via the Halpern-Bruno model, originally developed for protein-coding sequence [9]. This model incorporates selection on top of a mutational model. Essentially, the rate of evolution is modelled as a product of two terms: a site-independent rate of mutation and a site-dependent rate of fixation (*R*_*αβp*_ α *r*_*αβ*_*F*_*αβp*_ where *r*_*αβ*_ is the rate of mutation from *β* to *α* and *F*_*αβp*_ is the probability of fixation from *β* to *α* at position *p*. They use Kimura’s weak-mutation model (time of fixation ≪ time between fixations) to estimate *F* in terms of selection coefficients, which in turn is estimated from the equilibrium frequency (stationary distribution), which they call *f*, at position *p*. A key assumption is that *f* (which indicates the long-term distribution of nucleotides at a given locus over the course of evolution) is given by the corresponding column of the position weight matrix (which gives the distribution of nucleotides at that position across TFBS instances in a single species).

We make the contrary assumption, that the stationary distribution at a position is *not necessarily* given by the nucleotide counts for TFBS instances within a species. Our goal is to ascertain this distribution at each position within a TFBS. To do this within the Halpern-Bruno framework would require an independent way of estimating *F*, and it is not obvious how to do this.

We therefore opt for the simpler F81 and HKY85 models to ascertain stationary distributions. We find that position-specific stationary vectors (PSSVs) at TFBS loci reflect position weight matrices (PWMs), under both F81 and HKY85, but are weaker than the PWMs in terms of information content, and weaken further when looking at a larger group of mammals compared to using primates alone. Since we have no data on binding affinity in other primates or mammals, and are going purely by gapless alignments, this observation may arise from evolutionary divergence or turnover of TFBS. This varies by TF, with GABPA being strongly conserved among mammals, TCF12 being significantly weakened.

Focussing on CTCF, we find that PSSVs inferred for our approach are strongest for genuine CTCF motif instances in CTCF ChIP-seq peak regions (information loss *<* 10%); weaker for scrambled CTCF motifs in CTCF ChIP-seq peak regions, and for genuine CTCF motif instances in random genomic regions (information loss ≈ 15%–18%); and weakest for scrambled CTCF motifs in random genomic regions (information loss ≈ 23%-25%). It is known that there are frequently secondary motifs and sequence features in the neighbourhood of the TF core motif [25, 26], so it is plausible that an extended region around the core motif, including chance instances of scrambled motifs, may be under greater selection in ChIP-seq regions compared to random genomic regions. For a truly neutrally evolving region, one expects that the PSSV would be almost flat, but estimates of what fraction of the human genome is functional or under selective constraint range from 7% [27] to, controversially, 80% [28], with many authors agreeing that 10%–15% is plausible (see [29] for a discussion). It is likely that some fraction of our CTCF motif instances in random genomic regions are functional binding sites, and some fraction of our random genomic regions are in fact under selection pressure.

TFBS turnover has been reported widely earlier, in human/mouse [30] and fruitfly [31, 32]. Recently Krieger *et al*. [33] studied the effect of TFBS sequence variation and its effect on TFBS binding in two closely-related yeast species, considering both motif variation and chromatin accessibility, and find that imprecise motifs are bound to a high level by TFs and, in many cases, binding localization in the two species is conserved despite sequence divergence, suggesting “fast and flexible evolution” of TFBS. The same is likely true in other organisms.

It is widely recognized that PWMs are not adequate representations of TFBS complexity, which contain significant positional correlations [18] and multiple efforts have been made to go beyond the PWM model [2, 3, 4, 5]. As a complementary approach, we show that positional dependency effects can be examined by conditioning PSSVs on specific ancestral nucleotides.

We consider the F81 and HKY85 models. The primary difference is that HKY85 accounts for differing transition and transversion rates, which is an important factor in neutral evolution and in general phylogenetic problems. However, both models can reproduce any arbitrary stationary vector, and we observe that both models result in similar stationary vectors (which arises from a combination of mutation and purifying selection, not from mutation alone). The F81 model is conceptually simpler and we demonstrate an algorithm that greatly improves calculation time on moderate-sized trees (≲ 10 leaves), enabling rapid notebook-style analyses of sets of aligned TFBS (this speedup is important because, in estimating branch lengths or stationary vectors via multivariate nonlinear optimization, the likelihood function is called repeatedly, including for numerical derivatives). We plan to perform detailed evolutionary analysis of TFBS, and of CTCF in particular, in future studies, and hope our tool and methods will be useful to others.

## Author contributions

**Conceptualization:** RS

**Data Curation:** PS

**Formal Analysis:** PS, RS

**Funding Acquisition:** RS

**Investigation:** PS, RS

**Methodology:** RS

**Project Administration:** RS

**Resources:** RS

**Software:** PS, RS

**Supervision:** RS

**Validation:** PS, RS

**Visualization:** RS

**Writing – Original Draft Preparation:** PS, RS

**Writing – Review & Editing:** PS, RS

## Data and code availability

Processed data for primate and mammal sites, and code in the form of Jupyter notebooks, are available at https://github.com/rsidd120/TFBS-PSSV and have been archived in the Zenodo repository doi:10.5281/zenodo.10417053

## Appendix

### Models of nucleotide evolution, stationary vectors

For a detailed review of models of nucleotide evolution, see, eg, Tavaré [34]. A brief summary follows.

The instantaneous rate of mutation of nucleotides from *α* to *β* (*α, β* ∈ A, C, G, T ≡ 1, 2, 3, 4) is given by a 4 *×* 4 rate matrix *R*_*βα*_, whose columns sum to zero: *R*_*αα*_ =−Σ_*β*_ *R*_*βα*_. The “transition matrix” *T*_*βα*_(*t*) is the probability, given an ancestral nucleotide *α* at a given locus at time *T*, of observing *β* at the same locus at a later time *T* + *t*, and is equal to exp(*Rt*). The equilibrium distribution of nucleotides at a locus is denoted by 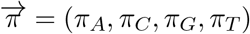 and, for a time-reversible model, we have

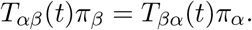

Here time *t* is geological time, measured in appropriate units. Transition matrices also satisfy the following multiplicative property

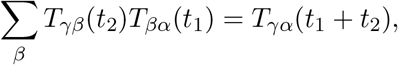

in other words, an intermediate node (here, the unknown *β*) with just one child can be removed from the tree and the distance of its child can be added to its distance from its parent. In combination with time-reversibility, this also implies that the branch lengths of the two children of a root node cannot be determined uniquely, only their sums can be, and indeed any node can be chosen to be the root without changing the likelihood (what Felsenstein called the “pulley principle” [11]).

Different models of nucleotide evolution differ in their choice of *R*. The simplest, the Jukes-Cantor model [7], assumes all off-diagonal elements of *R* are equal (*r*) and the diagonal elements are 3*r*. The Kimura model [8] distinguishes between transition rates (A ↔G, C ↔T, which are typically more frequent) and tranversion rates (all other mutations). Both of these result in a uniform equilibrium frequency 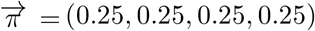.

Reproducing more general 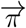 requires asymmetric matrices *R*. The simplest of these, the Felsenstein 1981 (F81) model [11], uses

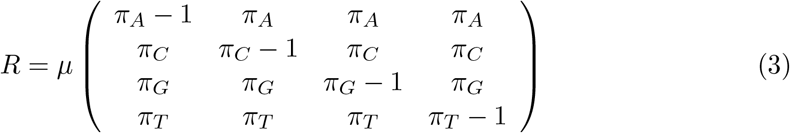

where *μ* is a mutation rate. This yields

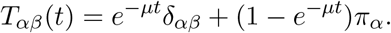

We choose units of time such that *μ* = 1, and rather than use the time *t*, we use a quantity *q* ≡ *e*^−*t*^ (which we previously called the “proximity” [13]):

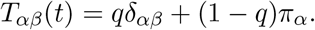

This has a simple interpretation: with a probability *q*, the nucleotide is unmutated from its ancestor, and with a probability 1 − *q*, it has undergone one or more mutation events, and also undergone selection, such that the probability of the new nucleotide *α* is given by the stationary distribution *π*_*α*_ indicating the relative fitness of *α*.

The Felsenstein 1984 (F84) [35] and the Hasegawa-Kishino-Yano 1985 (HKY85) [12] models generalize F81 to account for differing transition and transversion rates. We discuss HKY85 below, and its results on this problem are similar to F81. However, we demonstrate a fast likelihood algorithm for the F81 model (developing on an approach in a previous work by one of us [13]) which makes it more useful for this task. We argue that the similar results are because the evolution of TFBS are dominated by selection, so the differing rates of transition and transversion are not important: at most positions the rate matrix is dominated by the value of the equilibrium frequency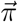 which indicates selection pressures at that position.

#### Likelihood calculation on a phylogenetic tree; Felsenstein’s algorithm

Consider a large collection of transcription factor binding sites in a species of interest, as well as orthologous sequence in other species. These species are leaves on a phylogenetic tree, assumed to be binary (each internal node has exactly two children). Two examples, featuring primates and mammals, are in figure 1. For a tree with *n* species (leaves), we label the leaves with numbers from 1 to *n* and the ancestral nodes with numbers from *n* + 1 to 2*n* − 1.

The branch lengths indicate the evolutionary distance. Rather than an additive distance in time units (*t*) we label the branches with multiplicative proximities *q* = *e*^−*t*^: in the F81 model, if node/leaf *i* has descended from its parent for a time *t*_*i*_ with a mutation rate *μ* (which we take to be 1), then 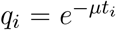, that is, *q*_*i*_ the probability that a given nucleotide under *i* is conserved (not mutated) from its parent. The proximities are therefore multiplicative along branches. For the root node, the children’s proximities are not independent and only their product can be determined (“pulley principle” mentioned above).

Each position within the binding site is a separate locus, with a separate collection of leaves, on the same tree. The likelihood, at a particular locus *x*, of seeing the collection of nucleotides {*S*^*k*^ } at the leaves (*k* denotes the leaf or species label) is then a product over the tree of all transition matrices along edges, summed over all ancestors. unknown ancestors. For the tree in fig. 1 (c), this is

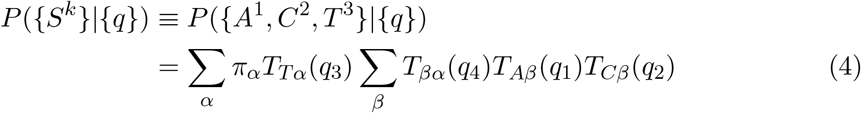

where, for the F81 model, as above, *T*_*βα*_(*q*) = *qδ*_*βα*_ + (1 − *q*)*π*_*β*_. For the HKY85 model too, explicit formulas for *T*_*βα*_ exist, discussed below. This sum depends only on the product *q*_3_*q*_4_ and not on their individual values. In our code we assign the right child of the root node a proximity *q* = 1, without loss of generality, and the root node also has a proximity 1 (it has no parent), so that for a tree of *n* leaves, and 2*n* – 1 nodes we need to calculate only 2*n* – 3 proximities.

Felsenstein, in the same paper where he introduced the F81 model [11], introduced a recursive algorithm to calculate this likelihood for any tree (and for any transition matrix), which can be summarised (for binary trees, but generalizable) as

- Define *L*_*i*_(*α*) = likelihood of leaves below node *i* given that the nucleotide at node *i* is *α*
- If *i* is a leaf node with nucleotide *x*, set *L*_*i*_(*α*) = *δ*_*αx*_
- Otherwise, let the two children of *i* be *j* and *k* with proximities *q*_*j*_ and *q*_*k*_. Then *L*_*i*_(*α*) = ∑*β T*_*βα*_(*q*_*j*_)*L*_*j*_(*β*) ∑ *γ T*_*γα*_(*q*_*k*_)*L*_*k*_(*γ*). (For non-binary trees this step can be generalized to a product over all children, with the appropriate number of sums.)
- Termination: let the root node be 2*n* − 1, then the likelihood of the leaves is ∑ _*α*_ *L*_2*n*−1_(*α*)*π*_*α*_

For efficiency, the values of *L*_*i*_(*α*) need to be stored, which we do within the tree structure.

This likelihood applies to nucleotides at one locus. It is assumed that nucleotides at different positions are independent, so the likelihood of a collection of sequences is the product of the likelihoods at individual loci. It is preferable to use log likelihoods to avoid underflows. There have been efforts to optimize the algorithm by sorting columns of the aligned sequence and storing and looking up identical calculations [36].

#### HKY85 model

The F81 assumes equal rates of transition and transversion. The HKY85 [12] model (and also the similar but non-equivalent F84 model [35]) are modifications of the F81 model to take account of a non-unity transition-transversion ratio *κ*. The rate matrix of the HKY model is

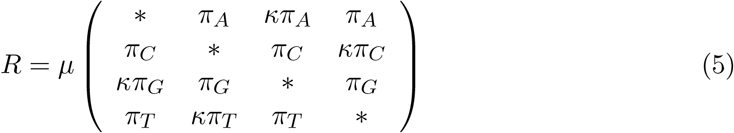

where the diagonal elements are such that columns sum to zero. We choose time units such that *μ* = 1 and, as before, define proximities as *q* = *e*^−*t*^ (however, these no longer have the simple interpretation of probability of conservation as in the F81 model).

We further define 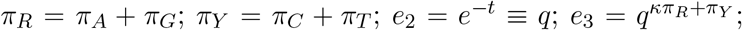 and 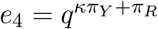.

In terms of these, the transition probabilities are: [12]

*Identity for purines* (*β* = *α* for *α* = *A, G*):

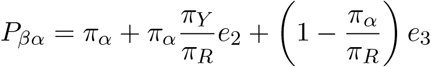

*Identity for pyrimidines* (*β* = *α* for *α* = *C, T*):

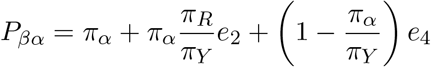

*Transition, purine* (*β* = *A, α* = *G* or vice versa):

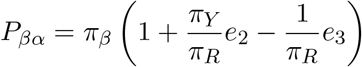

*Transition, pyrimidine* (*β* = *C, α* = *T* or vice versa)::

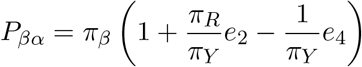

*Transversion* (all other cases):

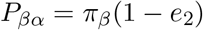

These can be derived by, for example, diagonalizing *R*, or via an argument involving likelihoods of different mutations analogous to the F81 picture [37].

